# Dispersal kernels influence the magnitude of environmental, biotic, and stochastic effects on the maintenance of metacommunity diversity

**DOI:** 10.1101/2025.08.12.669961

**Authors:** Nathan I. Wisnoski, Megan C. Szojka, Rachel M. Germain, Tadashi Fukami, Lauren G. Shoemaker

## Abstract

Dispersal plays a central role in shaping patterns of diversity in metacommunities. However, a primary focus on emigration rates may mischaracterize dispersal effects that actually arise from dispersal kernels. Kernels describe probabilistic movements between donor and recipient patches, but the influence of kernel shape on metacommunity diversity remains unclear. We used simulations to measure how kernels affect diversity across metacommunity scales and ecological contexts. We disentangled causes of these patterns using a novel approach quantifying the effects of environmental filtering, competition, stochasticity, and dispersal on fitness. Although metacommunities with shallow kernels followed expectations where emigration increased alpha-but decreased beta- and gamma-diversity, metacommunities with steeper kernels did not. Steeper kernels maintained regional diversity by reducing interspecific competition and stochastic extinctions, with dispersal conferring weaker benefits but less homogenization. Our work suggests dispersal kernels and emigration rates jointly regulate exposure to environmental variation and the balance of assembly mechanisms in metacommunities.

## INTRODUCTION

Metacommunity theory predicts that biodiversity at multiple spatial scales is shaped by dispersal across the landscape, with metacommunity models focusing on dispersal rates and their interactive effects with local patch conditions (Leibold & Chase 2018; Vellend 2016). Low-to-intermediate rates of dispersal are predicted to maximize local (α) diversity when biotic interactions are not stabilizing, such as when intraspecific competition is less than or equal to interspecific competition (Mouquet & Loreau 2003). In cases where competition is stabilizing (intraspecific > interspecific), mean α-diversity can be maintained at relatively higher dispersal rates (Thompson *et al*. 2020). High dispersal tends to spatially homogenize metacommunities, decreasing among-site (β) and regional (γ) diversity. Empirical support tends to be stronger for the β-diversity relationship than the α-diversity relationship (Cadotte 2006; Grainger & Gilbert 2016), a deviation from theory that might be explained by additional features of dispersal often overlooked in metacommunity theory. One consideration is that dispersal is a multi-step process that integrates emigration, movement, and establishment in a new location (Clobert *et al*. 2009). While metacommunity theory has focused primarily on the rate of emigration and its interactive effects with abiotic and biotic factors that influence establishment, the movement phase has received far less attention despite being the link between donor and recipient habitat patches.

The movement phase of dispersal is probabilistic because random features of individual organisms and the environment create variation in dispersal distances. The probability an individual moves a specified distance away from the source patch is described by a draw from a dispersal kernel; the full distribution of the kernel describes the expected distances, and corresponding probabilities, of dispersal for the full population (Levin *et al*. 2003; Rogers *et al*. 2019; Shoemaker *et al*. 2020). The dispersal kernel thereby influences patch accessibility to dispersers. Foundational metacommunity models often assumed spatially implicit dynamics where dispersal kernels were flat and dispersal was unconstrained by distance (Hubbell 2001; Mouquet & Loreau 2003). In contrast, spatially explicit models make dispersal probability a function of distance between patches. Short-distance movements typically occur more frequently than long-distance movements, but kernel shapes can vary with dispersal mode, environment, and biotic interactions (Beckman & Sullivan 2023; Bullock *et al*. 2017; Morales & Carlo 2006). Spatially explicit metacommunity theory has employed a range of dispersal kernels, including steep kernels with nearest-neighbor movements (Fournier *et al*. 2017) and gradually decaying kernels with regional-scale movements (Khattar & Peres-Neto 2024; Thompson *et al*. 2020). The effects of varying emigration rates have been well characterized for a subset of kernel shapes assumed in key models (Leibold & Chase 2018), but a joint consideration of emigration rates and kernel shapes is needed to understand their distinct and interacting effects on metacommunities, thereby breaking down the role of dispersal into its primary components.

Dispersal can generate unintuitive patterns in diversity because it has the potential to decouple abundances from population fitness. Fitness in a new patch may be low if the new environment is too far from a species’ niche optimum, if biotic interactions are too strong to allow growth, or if colonizing populations are too small to overcome the risk of stochastic extinction. If the demographic benefits of immigration outweigh the sum of local losses due to abiotic, biotic, or stochastic processes, then dispersal can support sink populations in suboptimal habitats where fitness is otherwise negative (Craig *et al*. 2025; Mouquet & Loreau 2003; Shmida & Ellner 1984). Therefore, dispersal can influence diversity by modifying demographic connectivity in relation to spatial and temporal environmental variation in the landscape, shifting the relative importance of community assembly processes, including environmental filtering, species interactions, and stochasticity. Spatially explicit simulations showed that sparse connectivity and spatial autocorrelation can generate clusters of connected patches with similar environmental conditions, which favors environmental filtering and reduces sink populations (Suzuki & Economo 2021). Linking metacommunity processes to their impacts on population fitness can reveal why populations exhibit non-negative long-term average fitness and are therefore predicted to persist over time.

Here, we examine the interactive effects of emigration rates and dispersal kernels on biodiversity in metacommunities and mechanistically link changes in diversity to the fitness consequences of environmental filtering, competition, stochasticity, and dispersal under different environmental scenarios. We consider a range of environmental scenarios because the pattern of spatial and temporal variation could alter the dispersal strategy (emigration rate and kernel shape) that is optimal for long-term regional persistence of the most species. For example, in environments with high environmental heterogeneity but low spatial autocorrelation, diversity could be best maintained if species send many dispersers broadly across the landscape (high emigration rates, shallow kernels). In contrast, in environments with high spatial autocorrelation and strong environmental gradients, diversity could be highest if fewer dispersers remain nearby (low emigration rates, steep kernels). In addition to habitat tracking across the landscape, dispersal is important in recolonizing local communities that experience species losses due to disturbances (Mausbach & Dzialowski 2019; Miller *et al*. 2011), so we also explore how our inferences from deterministic environments shift when stochastic patch-level disturbances influence dynamics.

We asked two main questions. Q1: How does the relationship between diversity and dispersal depend on emigration rate, dispersal kernel shape, and environmental variation? Q2: How do emigration rates and kernel shapes regulate the effects of dispersal relative to environmental filtering, biotic interactions, and stochastic processes, as assessed through their contributions to population fitness? We evaluated these questions using simulation models of metacommunities with systematically varied emigration rates and dispersal kernels in landscapes spanning environmental conditions (spatial and temporal heterogeneity) and disturbance regimes (Fig. 1). We linked model output to our motivating questions by quantifying diversity at different scales (Q1) and measuring population fitness and the demographic effects that led to species persistence in the metacommunities (Q2).

**Figure 1.**
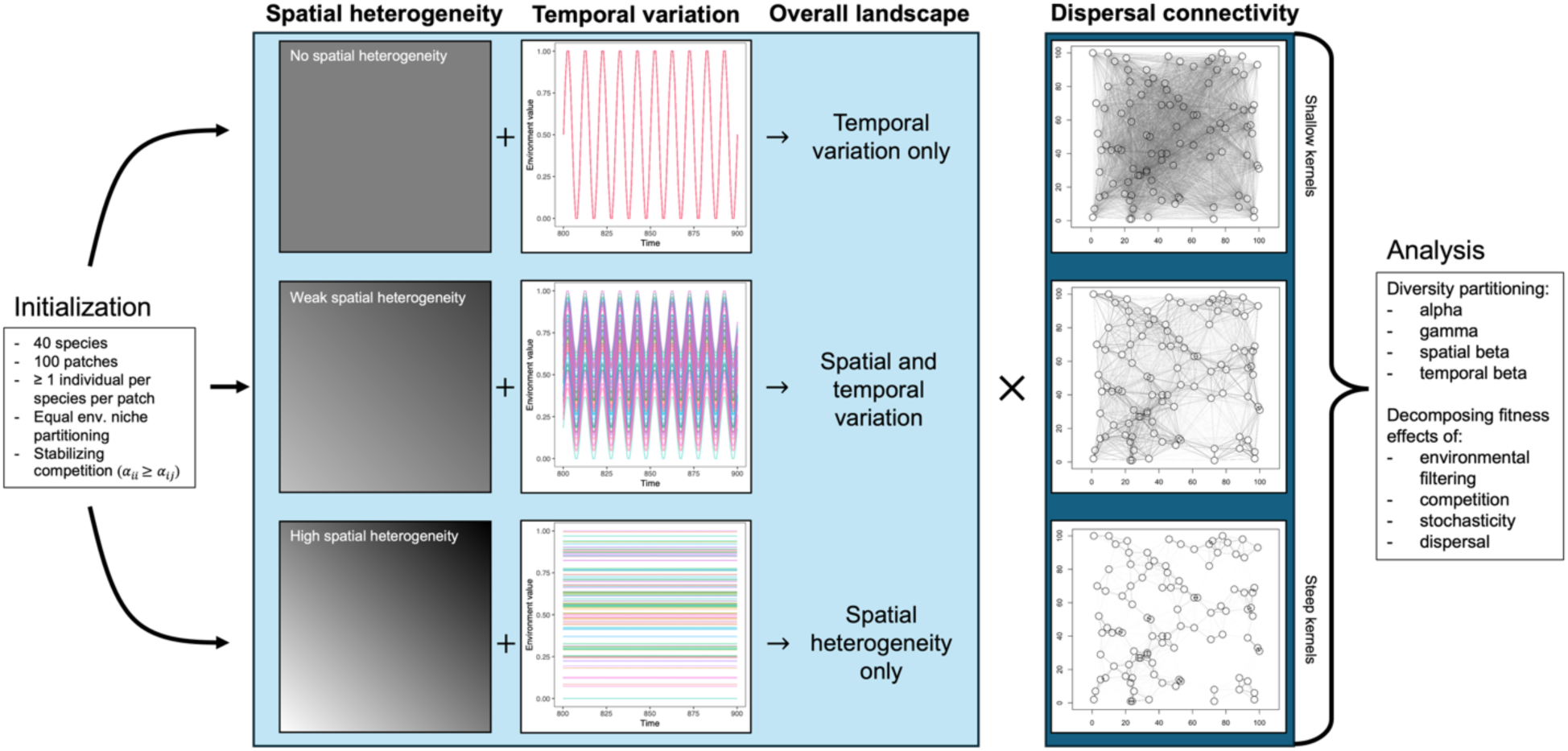
Overview of metacommunity model. Metacommunities were simulated in three landscape scenarios: temporal variation only, spatial and temporal variation, and spatial heterogeneity only. These scenarios arose from the addition of temporal variation to differing levels of background spatial heterogeneity and were crossed with different levels of dispersal connectivity. The level of connectivity due to dispersal was determined by the intersection of the spatial location on the landscape and the dispersal kernel shapes of the constituent species in the metacommunity. We analyzed metacommunity diversity along gradients of emigration rate and dispersal kernel steepness and decomposed fitness into the effects of community assembly processes.

## METHODS

To examine the effects of dispersal kernels on metacommunity diversity, we simulated metacommunities with varying dispersal properties and spatiotemporal patterns of environmental variability. We then analyzed the long-term diversity and fitness patterns in simulated dynamics (averaged over the final 100 timesteps) following an initial 1000-timestep burn-in period that removed transient dynamics.

### Metacommunity model overview

We modeled metacommunities using a process-based framework focused on abiotic responses, density-dependent biotic interactions, stochasticity, and spatially explicit dispersal (Shoemaker & Melbourne 2016; Thompson *et al*. 2020; Wisnoski & Shoemaker 2022). Change in population size of species *i* in patch *x* (*N*_*ix*_) depends on local growth (*G*_*ix*_) and the balance between emigration (*E*_*ix*_) and immigration (*I*_*ix*_)

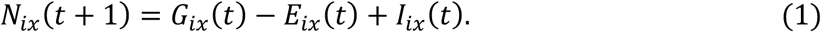

Local recruitment of species *i* in patch *x*, in turn, depends on several local-scale processes: abiotic conditions, biotic interactions, and demographic stochasticity. The abiotic effect is modeled through *r*_*ix*_(*t*), i.e., the density-independent per capita growth rate of species *i* in patch *x* at a given point in time in the absence of any biotic interactions or stochasticity. *r*_*ix*_(*t*) is a Gaussian function of the environmental condition (*η*_*x*_(*t*))

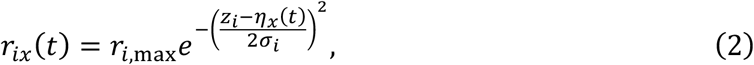

centered on the species’ niche optimum (*z*_*i*_) with standard deviation equal to its niche breadth (*σ*_*i*_). In other words, *r*_*ix*_(*t*) defines the range of environments that are suitable (*r*_*ix*_ ≥ 1) vs unsuitable (*r*_*ix*_ < 1). Density-dependent biotic interactions were modeled by multiplying per-capita interaction coefficients (*α*_*ij*_; restricted to be competitive such that *α*_*ij*_ > 0) by the number of neighbors of species *j*, *N*_*j*_ in equation 3. Incorporating both abiotic and biotic drivers, local dynamics for each patch *x* follow the discrete-time Beverton-Holt model (Beverton & Holt 1957). To incorporate demographic stochasticity, the final population growth of a species in a patch, *G*_*ix*_(*t*), was drawn from a Poisson distribution (Shoemaker & Melbourne 2016):

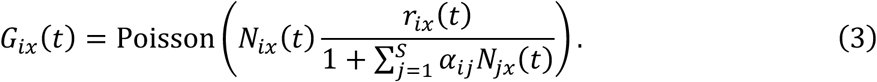

Dispersal in equation 1 was modeled with separate emigration, *E*_*ix*_(*t*), and immigration, *I*_*ix*_(*t*), terms. Emigrants of species *i* were randomly selected to leave patch *x* following a binomial distribution, *E*_*ix*_(*t*)∼Binomial(*n* = *N*_*ix*_(*t*), *p* = *d*), where *d* is the per capita probability of emigration. Therefore, 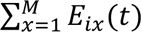 quantifies the total number of emigrants across all *M* patches in the metacommunity that get redistributed via dispersal to new patches.

Immigration was determined using a negative exponential dispersal kernel

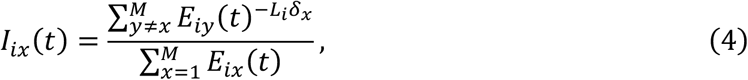

where *δ*_*x*_ is the distance between patches on the landscape and *L*_*i*_ regulates the shape of the dispersal kernel. When *L*_*i*_ = 0, the metacommunity is spatially implicit with global connectivity. As *L*_*i*_ increases, the probability of successful dispersal decreases more rapidly with distance because the dispersal kernel gets steeper. In this analysis, we vary emigration rate (*d*) and dispersal kernel exponent (*L*_*i*_) to investigate their contributions to diversity.

### Environmental variation

We modeled three general environmental scenarios along a gradient from temporal variation only, to both spatial and temporal variation, to spatial variation only (Fig. 1). To generate these different environmental scenarios, we first created a spatially explicit landscape in two dimensions (*x*_1_, *x*_2_): 100 × 100 units. Across the landscape, we generated spatially autocorrelated mean environmental conditions of a single abiotic variable 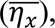 which increased linearly in both dimensions, 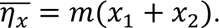 We varied the steepness (*m*) of the environmental gradients from *m* = 0 (no spatial heterogeneity) to *m* = 0.1 (weak heterogeneity), to *m* = 1000 (high heterogeneity) in each dimension, which described the mean environmental conditions for each patch. We generated temporal variability around these mean conditions by simulating a regional abiotic climate time series using a sine wave with 10-year cycles. The amplitude (*A*) of the sine wave was 5 climate units with angular frequency, *ω* = 2*π*/100. For each location in the landscape grid, we added this climate time series to the temporal mean condition *η̅*, generating temporal dynamics for each patch *x* in the landscape: 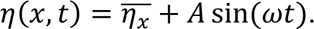

To align the environmental range with the range of species niche optima, we rescaled the spatiotemporal series to the range [0,1]. The rescaling made variability purely temporal when *m* = 0 (because there is no spatial heterogeneity), both spatially and temporally heterogeneous with patches shifting in and out of species’ abiotic niche limits when *m* = 0.1, and approximately purely spatial when *m* = 1000 (which overwhelmed any biologically relevant temporal variation on the order of ±5 units after rescaling). After normalization, the environmental conditions for each patch, *η*(*x*, *t*), were spatially and temporally autocorrelated, where patches have different mean conditions but experience synchronous regional-scale climatic fluctuations (Fig. 1).

We simulated the above three scenarios both without and with disturbances. We introduced stochastic local, patch-level disturbance events to the environment, which generated an additional unpredictable spatial and temporal level of variation across the landscape. Disturbances led to the removal of all local biomass, with all species going temporarily locally extinct. Local disturbances were binomially distributed with a per patch extinction probability of 0.01 per time. With 100 patches and 100 years in the final simulations, each patch was disturbed, on average, once during the analyzed time window, and an average of one patch was disturbed in the metacommunity per time step. As each disturbance event occurred independently of each other, environmental stochasticity was not spatially or temporally autocorrelated in the model.

### Simulation procedure

We initialized metacommunities with 40 species distributed across 100 patches that were placed on the 100 × 100 landscape by sampling locations uniformly at random. The random placement means that some patches will be more connected than others, and spatial structure is not symmetric as it would be in a full lattice grid. A single individual of each species was introduced to each patch to eliminate dispersal limitation in initial colonization. Additional individuals from each species were then randomly introduced to each patch with a probability of Poisson(*λ* = 0.5) per species per patch at time steps [10, 20, ⋯, 200] to introduce stochastic elements to the colonization process equivalent to dispersal from a mainland into the metacommunity (Lu 2021; Overcast *et al*. 2023; Thompson *et al*. 2020). We simulated metacommunities with a burn-in period of 1000 timesteps to remove any potential effects of initial conditions and analyzed the following 100 timesteps of the simulation, which correspond to the final ten 10-year environmental cycles.

The 40 species had niche optima evenly distributed along the rescaled environmental gradient [0,1] and all had maximum growth rates *r*_*i*,max_of 5 yr^-1^. We varied the strength of intraspecific competition (*α*_*ii*_) relative to interspecific competition (*α*_*ij*_), simulating scenarios of stabilizing competition (*α*_*ii*_ ≥ *α*_*ij*_) by assigning competition coefficients randomly following a uniform distribution from the range [0, *α*_*ii*_]. In this scenario, species in the metacommunity may have no interaction or interact less than or as strongly as intraspecific competitive interactions, which were fixed for all species (*α*_*ii*_ = 0.05). We simulated 20 emigration rates, *d*, in the range [10^−5^, 1], distributed evenly in logarithmic (base 10) space. We compared 10 dispersal kernel shape parameters, *L*_*i*_, for the negative exponential distribution distributed evenly in logarithmic space in the range [10^−4^, 1], plus the boundary condition of *L*_*i*_ = 0. If *L*_*i*_ = 0, the kernel is flat and connectivity is global; if *L*_*i*_ = 1, then connectivity is localized to only nearby neighboring patches (Fig. 1). We simulated 10 replicate landscapes (each with a different spatial configuration, which altered connectivity and the underlying distribution along the environmental gradients) to detect strong patterns that emerge across relationships that may rely on specific landscape types. On each landscape we ran simulations of every parameter combination (emigration rate, kernel shape, environmental scenario, and disturbance), for a total of 12,000 simulations.

### Analysis of Model Dynamics

#### Patterns of α-, β-, and γ-diversity

All analyses focused on the last 100 timesteps of the simulated 100-patch metacommunities. To analyze the effects of dispersal kernels on biodiversity in the metacommunity, we partitioned diversity across scales and along spatial and temporal dimensions (Fig. 1). We partitioned diversity by computing metacommunity γ-diversity and mean patch-level α-diversity at each time point and averaging over time. To compute β-diversity, we used variance-based approaches to have a comparable metric for spatial and temporal turnover (Legendre & De Cáceres 2013). For each simulation, we computed spatial β-diversity (across all 100 patches), averaged through time, and temporal β-diversity of each patch (over 100 time points), averaged across all patches.

#### Partitioning the fitness effects of metacommunity processes

We developed a novel approach to characterize how the dispersal kernel regulated the influence of key community assembly mechanisms—environmental filtering, biotic interactions, stochasticity, and net dispersal—on species abundances and distributions (HilleRisLambers *et al*. 2012). To do so, we simulated expected fitness when including versus excluding different processes that regulate metacommunity dynamics. By comparing expected fitness and assessing population sizes with versus without each component, we quantified the per capita effects of each mechanism on mean fitness (local per capita growth). This approach allowed us to assess the fitness benefits and costs associated with each process regulating metacommunity dynamics. By looking at a finer resolution than typically considered in metacommunity studies, we aimed to better link large-scale patterns of diversity with demographic changes of populations in each patch.

First, we assessed environmental filtering by quantifying the per capita effects of environmental mismatch. To do so, we quantified the reduction in per capita density-independent growth due solely to environmental effects

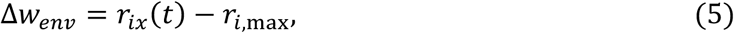

and then took the average across all 40 species, weighted by abundance, in all 100 patches over the final 100 timesteps. This quantified the per capita demographic cost of being in a mismatched environment at the metacommunity scale. The maximum value for this effect is 0, which would be the case if all species occupied only environments that directly matched their niche optima. Negative fitness effects indicate greater costs of environmental filtering operating across the metacommunity.

Next, we quantified competition by summing the per capita effects of all individuals in local communities on the focal species’ per capita growth. For species *i* in patch *x*, we quantified total intraspecific competition as *α*_*ii*_*N*_*ix*_ and interspecific competition as ∑*α*_*ij*_*N*_*jx*_ for all *i* ≠ *j*. We then re-computed the Beverton-Holt growth equation based on these different denominators and compared recruitment with only intraspecific or interspecific competition to the expectation in the absence of competition (i.e., exclusively environmental filtering operating). To measure the fitness cost of competition, we calculated the difference in per capita recruitment between the expectation with competition (intraspecific only or interspecific only) and the expectation in the absence of competition (i.e., only environmental effects), normalized to the average abundance of each species in each patch averaged over the 100 timesteps:

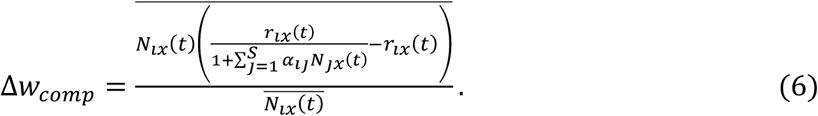

Note that the fitness effects of competition are also negative, indicating the additional reduction in per capita growth rates due to competitive interactions after accounting for environmental filtering.

To assess the impact of demographic stochasticity, we quantified the number of local extinctions due to demographic stochasticity, on average, per metacommunity. To infer a stochastic extinction, we counted the number of times we observed a population predicted to have positive local growth based on the deterministic, patch-level expectation (Eq. 3, 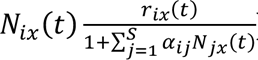) but had a local population size of zero after accounting for demographic stochasticity, given population sizes were drawn from a Poisson distribution. This pattern would indicate a local extinction due to demographic stochasticity after accounting for abiotic and biotic effects. We summed the number of extinctions for each metacommunity and then averaged across replicate metacommunities.

We quantified the fitness effects of dispersal on population dynamics by comparing per capita growth estimates before and after dispersal. For some populations, dispersal had a net negative effect because excess individuals were redistributed to other populations and not replaced (local cost of emigration). For other populations, dispersal had a net positive effect because they received more immigrants than the number of emigrants lost. We quantified the overall long-term impact of dispersal on population fitness in the metacommunity as the net population change due to immigration and emigration expressed on a per capita basis for the local population and then computed a weighted average across all patches and species and timesteps. Although many populations exhibited negative fitness effects of dispersal at the patch level for some portion of time, the average fitness effects were non-negative because species with negative long-term fitness were gradually eliminated over time. We focus on the metacommunity averages here because these quantities relate to species persistence and the role of dispersal in the maintenance of regional biodiversity.

## RESULTS

### Patterns of alpha-, beta-, and gamma-diversity

Mean α-diversity was maximized at intermediate-to-high emigration rates except at the highest dispersal kernel exponents (Fig. 2A). As kernel exponents increased (i.e., kernels became steeper with more localized dispersal), higher emigration rates were necessary to maintain alpha diversity. Increases in the dispersal kernel exponent yielded a transition in the relationships between mean alpha diversity and emigration rate: from humped-shape relationships (with highest α-diversity at moderately high emigration rates) to monotonic relationships (with highest α-diversity at high emigration rates). Across environmental scenarios, higher spatial variation tended to allow for higher mean alpha diversity. The addition of local disturbances had minor effects on α-diversity, but led to a decrease in α-diversity for the steepest dispersal kernels and at the lowest emigration rates.

**Figure 2.**
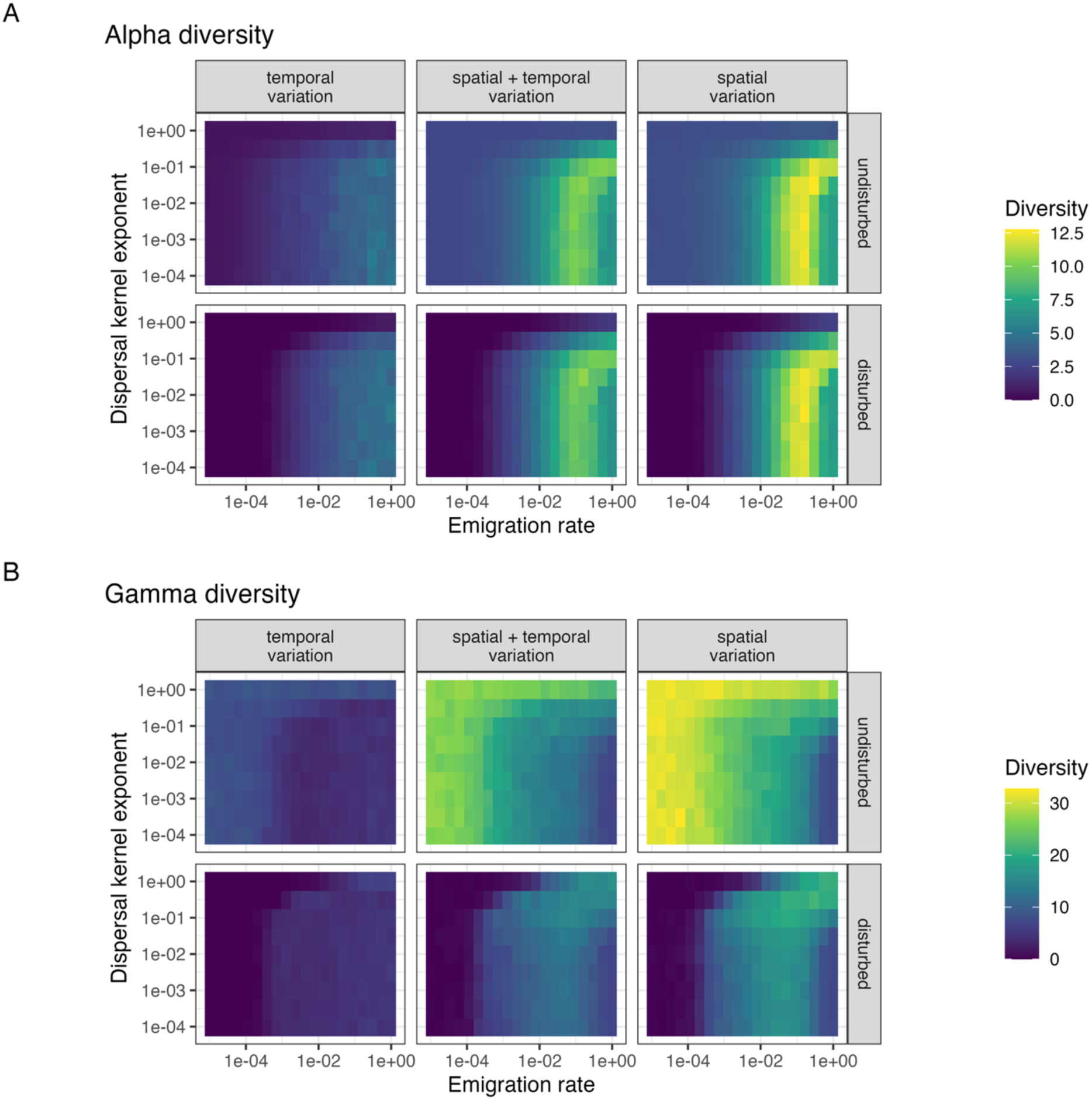
Local α- and regional γ-diversity along gradients of emigration rates and dispersal kernel exponents (i.e., steepness). (A) Mean α-diversity for each parameter combination in undisturbed and disturbed scenarios across three types of environmental variation: temporal variation in a spatially homogeneous landscape, spatial and temporal variation, and purely spatial variation in a temporally static landscape. (B) Mean γ-diversity for each parameter combination. Note that the heatmaps display means across 10 replicate simulations per parameter combination, summarizing 12,000 total simulations, and that the color scales differ between panels A and B to enhance comparability between disturbed and undisturbed scenarios and to account for different magnitudes between the two metrics.

At the metacommunity scale, the effects of dispersal on γ-diversity were dependent on spatial heterogeneity and disturbance (Fig. 2B). High spatial heterogeneity allowed the maintenance of about 4 times higher γ-diversity (*γ̅* = 23.6) than landscapes with temporal variation alone (*γ̅* = 5.77). In the absence of disturbance, γ-diversity was eroded by high emigration rates and flat kernels (i.e. with smaller exponents). As dispersal kernels became steeper (i.e., exponents approached 1), higher emigration rates were able to maintain regional diversity that would otherwise have been lost with flatter kernels. Comparable values of γ-diversity were obtainable under either high emigration/steep kernel or low emigration/shallow kernel scenarios. In contrast, when disturbances were imposed, γ-diversity was maximized by a combination of high emigration rates with steep kernels. Disturbances generally reduced γ-diversity and introduced a transition in γ-diversity being a unimodal function of emigration rate for flat dispersal kernels and a monotonically increasing function of emigration rate for steeper kernels.

The effects of dispersal on turnover, or β-diversity, were also dependent on the form of environmental variation (Fig. 3). Average spatial β-diversity was higher than the average temporal β-diversity, and β-diversity typically increased with spatial heterogeneity. In the absence of disturbance, spatial β-diversity generally declined with higher emigration rates and flatter kernels in the absence of disturbance (Fig. 3A). Local disturbances made spatial β-diversity a unimodal, hump-shaped function of emigration rate in shallow-to-moderate dispersal kernels, but a monotonically increasing function of emigration rate as kernels became steeper. Temporal β-diversity was highest at intermediate emigration rates and kernel exponents, particularly in disturbed landscapes (Fig. 3B). Kernel shape typically had a weaker effect on temporal β-diversity than emigration rate or environmental variability. In the absence of disturbance, local composition had relatively low temporal variability.

**Figure 3.**
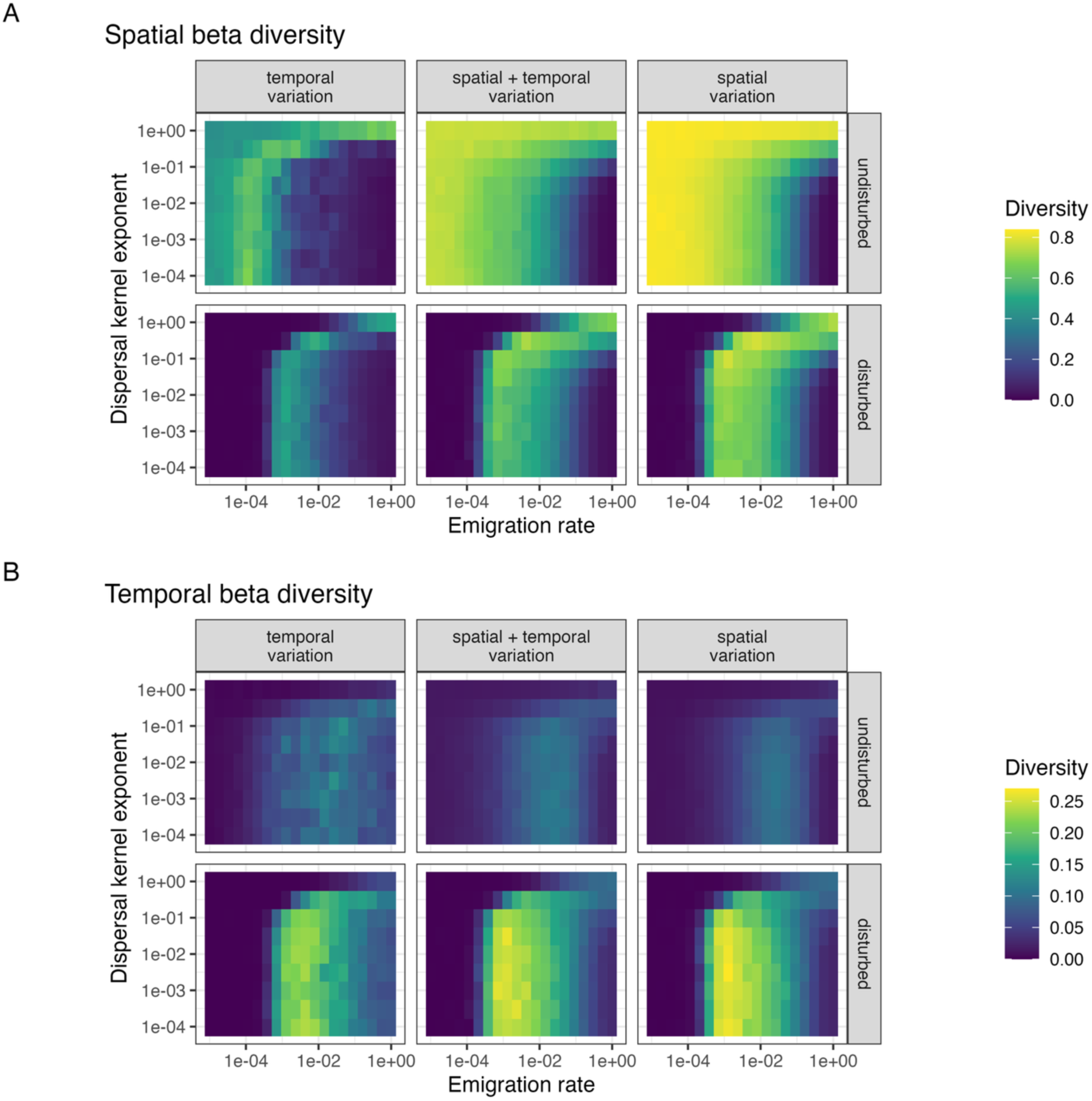
Spatial and temporal β-diversity along gradients of emigration rates and dispersal kernel exponents (i.e., steepness). (A) Mean spatial β-diversity for each parameter combination in undisturbed and disturbed scenarios across three types of environmental variation: temporal variation in a spatially homogeneous landscape, spatial and temporal variation, and purely spatial variation in a temporally static landscape. (B) Mean temporal β-diversity for each parameter combination. Note that the heatmaps display means across 10 replicate simulations per parameter combination, summarizing 12,000 total simulations, and that the color scales differ between panels A and B to enhance comparability between disturbed and undisturbed scenarios and to account for different magnitudes between the two metrics.

### Partitioning the fitness effects of metacommunity processes

Fitness costs due to mismatches between environmental conditions and species occurrence varied with dispersal kernel shape, emigration rate, and patterns of environmental variation (Fig. 4). Shown in the graphs are the scenarios without disturbance to allow calculation of fitness effects before environmental stochasticity (which was density-independent). With purely temporal variation, all patches experienced the same conditions when averaged through time, and for most of the environmental cycle species encountered suboptimal conditions, which was exacerbated for species with niche optima towards the extreme ends of the environmental range. Consequently, fitness effects of the environment were negative (Fig. 4A). However, when the environment had spatial structure, fitness costs of the environment could be reduced (becoming less negative) via dispersal. Generally, steeper dispersal kernels (i.e., higher exponents, lighter colors) with lower emigration rates experienced lower fitness costs due to environmental/occurrence mismatches, while shallow kernels (i.e., lower exponents, darker colors) with intermediate-to-high emigration rates had higher costs, where the extent of costs depended on emigration rate. With shallow kernels (darker colors), costs were highest at intermediate-to-high emigration rates, with a slight reduction in costs at the highest emigration rates as patches became homogenized. Steeper kernels (lighter colors) always experienced a decrease in mean fitness with increased emigration rate.

**Figure 4.**
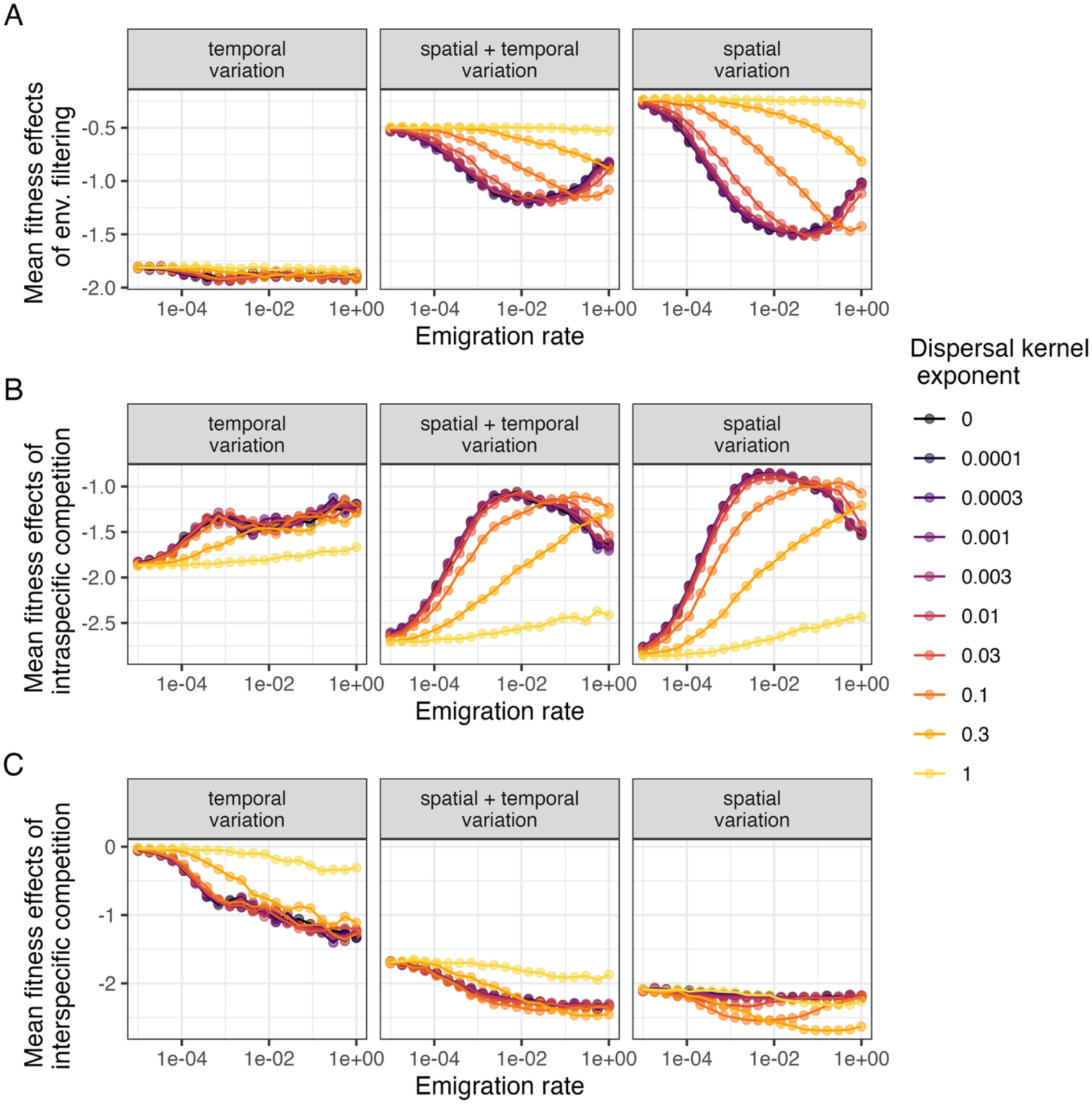
Environmental and biotic filtering vary with emigration rate and dispersal kernel. (A) Environmental filtering was measured as mean fitness consequences of environmental mismatch on all populations the metacommunity. (B) Fitness costs due to intraspecific competition averaged across all populations. (C) Fitness costs due to interspecific competition averaged across all populations. Note, because these metrics quantify costs, more negative values indicate stronger constraints on population growth due to the given process. Results are shown for environmental variation ranging from purely temporal, to spatial and temporal, to purely spatial. Each point is the mean value of the metric across simulations, while lines emphasize differences among dispersal kernels.

Dispersal kernel shape, emigration rate, and environmental variation jointly regulated the fitness costs of competition, with kernel exponent having weaker effects at low emigration rates (Fig. 4B-C). Without spatial heterogeneity, higher emigration rates and shallower kernels generally weakened intraspecific competition (Fig. 4B). With spatial heterogeneity, intraspecific competition was strongest at low emigration rates and weakened with increased emigration. Shallow kernels showed hump-shaped fitness responses to emigration (darker colors). In contrast, the negative effects of interspecific competition generally became stronger with higher emigration rates, except under the spatial-variation-only scenario (Fig. 4C). Interspecific competition was weaker without spatial heterogeneity.

Demographic stochasticity caused more local extinctions at intermediate-to-high emigration rates, and with shallower kernels (Fig. 5A). Shallow kernels exhibited a hump-shaped, unimodal pattern, where stochastic extinctions were maximized at intermediate emigration rates, while steeper kernels led to increasing numbers of local extinctions with increasing emigration rates. Extinctions due to demographic stochasticity were not driven directly by environmental filtering and competitive exclusion but by variation in the recruitment process. Fewer stochastic extinctions were observed in the absence of spatial heterogeneity.

**Figure 5.**
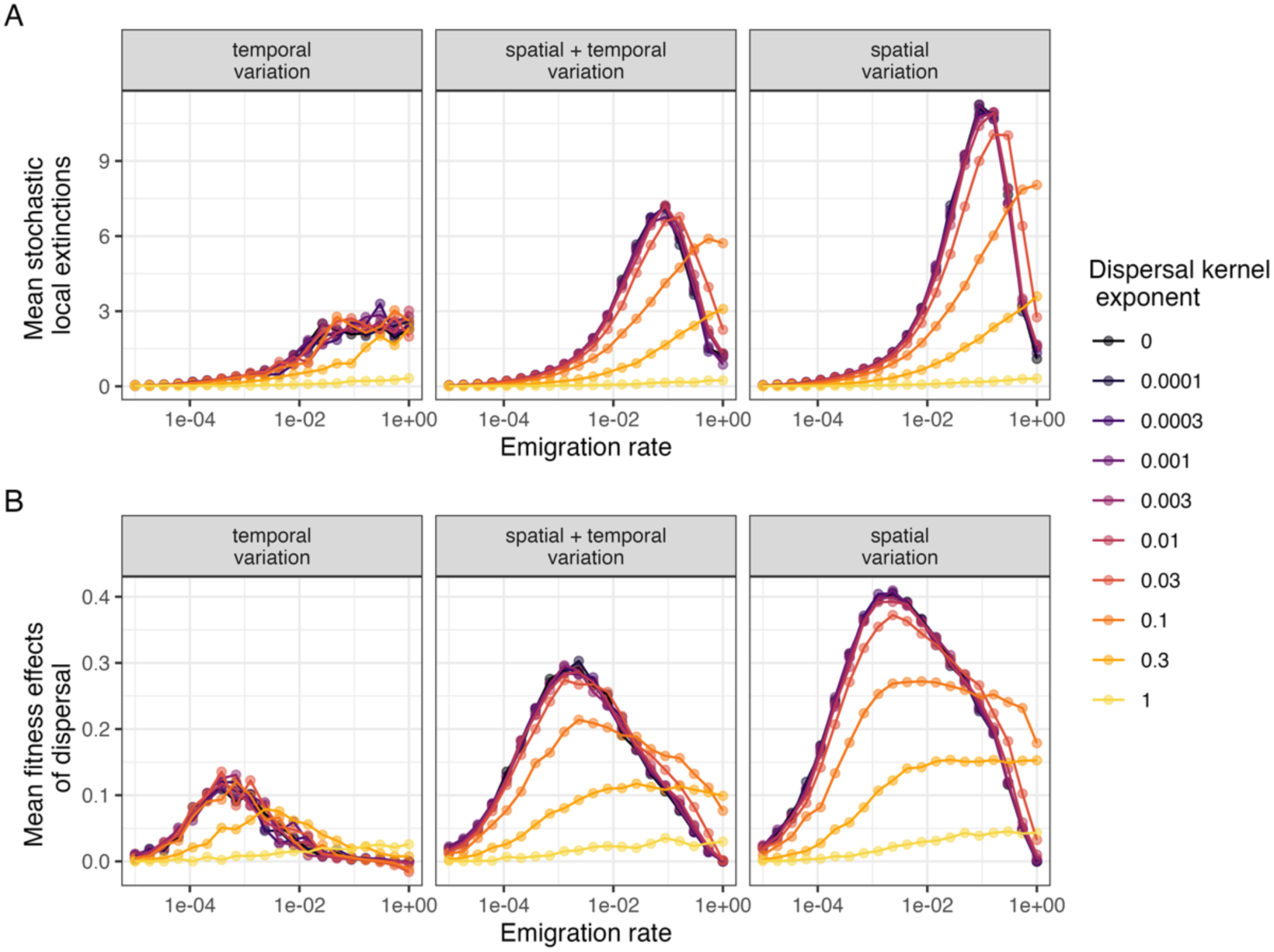
The population-level impacts of demographic stochasticity and dispersal along gradients of emigration rate and kernel shape. (A) A local extinction was due to demographic stochasticity if a population was predicted to have positive recruitment after environmental filtering and biotic interactions but went extinct due to demographic stochasticity. (B) Fitness benefits due to dispersal, after accounting for environmental filtering, biotic interactions, and demographic stochasticity. Dispersal effects quantify the average gain in fitness of each population in the metacommunity due only to dispersal (both immigration and emigration). Results are shown for environmental variation ranging from purely temporal, to spatial and temporal, to purely spatial. Each point is the mean value of the metric across simulations, while lines emphasize differences among dispersal kernels.

The average net effects of dispersal on fitness at the metacommunity scale (i.e., the fitness benefits of immigration minus the fitness costs of emigration) were generally positive, but the magnitude of effects varied with emigration rate, kernel shape, and environmental scenario (Fig. 5B). Dispersal provided the largest fitness benefits in spatially heterogeneous landscapes. The shape of the dispersal kernel generated wide variation in the fitness effects of dispersal across the emigration rate gradient. Typically, for shallow kernels (darker colors), the benefits of dispersal were maximized at low-to-intermediate emigration rates. For steeper kernels (lighter colors), the maximum benefits of dispersal were reduced, but dispersal benefits occurred over a wider range of emigration rates, with maxima at slightly higher emigration rates.

## DISCUSSION

By simultaneously and systematically varying both dispersal kernel shapes and emigration rates, our simulations quantified how these two distinct components of dispersal regulate diversity by mediating the relative importance of environmental, biotic, and stochastic effects on population fitness. Steeper dispersal kernels typically reduced α-diversity but preserved γ-diversity by preventing spatial homogenization (maintaining β-diversity), especially in spatially variable landscapes. Shallow kernels maintained similar γ-diversity but only if emigration rates were sufficiently low to prevent homogenization. Disturbances introduced minimum dispersal thresholds necessary for regional persistence, and steeper kernels with higher emigration rates could promote γ-diversity better than shallow kernels with lower emigration rates. Consistent with existing theory, shallow kernels generally conferred higher fitness benefits at low-to-intermediate emigration rates, but our results suggest that steeper kernels may increase the benefits of dispersal across a wider range of emigration rates. By decomposing patterns of diversity and analyzing the fitness effects at the population level, we were able to attribute the benefits of localized dispersal to the fact that steeper kernels reduced the costs of environmental mismatch, favored intraspecific over interspecific competition, minimized stochastic extinctions, and expanded the range of emigration rates at which dispersal increased fitness. By quantifying the demographic effects underlying emergent biodiversity patterns, our work demonstrates the interplay between different aspects of the dispersal process in regulating the balance between environmental, biotic, and stochastic effects in metacommunities.

Regional persistence is facilitated by dispersal kernels that align with the scale of environmental heterogeneity. Temporal fluctuations in the absence of spatial heterogeneity did not allow species to spatially partition the landscape based on environmental favorability; instead, spatial variation was necessary for habitat partitioning in abiotic dimensions. Metacommunities with shallow kernels had species, on average, more displaced from their environmental optima (Fig. 4A), indicating a mismatch between the scales of dispersal and environmental variation, which led to higher fitness costs of environmental mismatch. The combination of shallow dispersal kernels and intermediate emigration rates led to the highest mean α-diversity, despite the dispersal-environment scale mismatch (Fig. 2A). Intermediate rates of longer distance dispersal provided the biggest per capita benefits of dispersal at the metacommunity scale (Fig. 5B), as local costs of environmental mismatch were offset by fitness benefits of efficient environmental tracking in other patches. Recent empirical studies have shown that dispersal from productive patches can maintain sink populations in environmentally suboptimal patches, emphasizing the regional environmental context that allows sink populations to depend on productive source patches (Craig *et al*. 2025). Our models show that these source-sink dynamics may be undermined by higher emigration rates, which can increase interspecific competition (Fig. 4C), decrease diversity at the local scale, and lead to spatial homogenization (Figs. 2-3). Flatter kernels facilitated spatial homogenization, which eroded metacommunity diversity and reverted the metacommunity to regulation primarily by intraspecific competition (Fig. 4B). These dynamics provide additional mechanistic insight into foundational mass effects metacommunity models (Mouquet & Loreau 2003).

In contrast, steep kernels provide a mechanism of retaining local dominance and promoting regional diversity and stability in less dynamic landscapes. Consistent with species sorting metacommunity models, species with steep kernels mainly persisted in optimal patches, and fitness costs due to environmental mismatch were minimal, except under high emigration rates (Fig. 4A). Because populations were well matched to the environment, they had fewer stochastic extinctions, although they still benefited from dispersal (Fig. 5). Steeper dispersal kernels restrict movement and concentrate intraspecific interactions within patches, a stabilizing spatial coexistence mechanism due to its tendency to promote self-limitation (Amarasekare 2003; Snyder & Chesson 2003, 2004). Other studies have shown that increased dispersal can undermine regional coexistence by weakening spatial coexistence mechanisms like the spatial storage effect and fitness-density covariance, which depends on spatial segregation of density into high-fitness patches (Shoemaker & Melbourne 2016). In our models, intraspecific competition was stronger than interspecific competition for all dispersal kernel shapes at low emigration rates due to local species monodominance, but steeper kernel shapes promoted intraspecific limitation and buffered interspecific competition. Consequently, steeper kernels may counteract environmental and competitive costs, and preserve regionally stabilizing spatial storage effects and fitness-density covariation.

Dispersal kernels and emigration differentially regulate the impact of environmental and demographic stochasticity on population persistence. Environmental stochasticity, modeled here in the form of random local disturbances, typically had detrimental effects on γ-diversity (Fig. 2B), but different dispersal strategies were able to preserve diversity even in the face of disturbances, such as by shifting the metacommunity towards a more locally dynamic system that maintained moderate γ-diversity (Fig. 3B). Demographic stochasticity is often considered a concern for small populations isolated by dispersal limitation (Lande 1993). However, our results suggest that local extinctions due to demographic stochasticity can also be caused by high rates of dispersal into unfavorable environments, or excess emigration out of favorable habitats, both of which decrease local population sizes (Fig. 5A). The interplay between dispersal and stochasticity also has implications for biological invasions by influencing the propagule pressure needed to surpass abundance thresholds imposed by stochastic limits on establishment (Simberloff 2009). Our work suggests that dispersal may expand species ranges into environments that could, in principle, support their growth, but is not enough to overcome the risk of stochastic extinctions. Building on previous metacommunity work highlighting the importance of stochastic processes in regulating biodiversity (Hubbell 2001; Leibold *et al*. 2004; Shoemaker *et al*. 2020; Vellend 2010), our study demonstrates potentially overlooked interactions between emigration, dispersal kernels, and stochasticity.

Our results are consistent with the notion that the environment promotes coexistence if it enables spatial niche differentiation. Spatial niche differences may be strengthened when the spatial scale of landscape heterogeneity is large relative to typical dispersal distances and when spatial structure persists for multiple generations (Snyder 2008). Scaled up to a larger number of species, our simulations showed regional diversity was highest with purely spatial heterogeneity and steep dispersal kernels. By restraining movements to occur within neighboring patches in a spatially and temporally autocorrelated landscape, species stayed near favorable conditions (Suzuki & Economo 2021). Consistent with species sorting, restricting species distributions minimized environmental mismatch and promoted intraspecific competition (Fig. 4). Spatial niches were undermined by reduced spatial heterogeneity or flattened dispersal kernels, which increased interspecific competition and environmental costs and decreased γ-diversity. In more spatiotemporally variable or patchy landscapes, the relative costs and benefits of dispersal kernels may differ, giving rise to bet-hedging strategies involving flatter kernels and long-distance dispersal. Nevertheless, our results are consistent with the hypothesis that the tension between homogenization and spatial differentiation is mediated by the scale of environmental variation (Zelnik *et al*. 2024). Our work shows that spatial differentiation and homogenization relate to interactions between dispersal kernels, emigration rates, and local abundances that influence the probabilistic nature of dispersal as a key driver of processes and patterns in metacommunities.

## ACKNOWLEDGMENTS

This project was paid for in part with federal funding (to NIW) from the U.S. Department of the Treasury, the Mississippi Department of Environmental Quality, and the Mississippi Based RESTORE Act Center of Excellence under the Resources and Ecosystems Sustainability, Tourist Opportunities, and Revived Economies of the Gulf Coast States Act of 2012 (RESTORE Act). LGS was supported by the National Science Foundation (NSF) grants 2019528 and 2033292. MCS was supported by a Natural Sciences and Engineering Research Council of Canada (Postgraduate Scholarships – Doctoral program Award), Award number 557373-2021 and NSF 2033292. The statements, findings, conclusions, and recommendations are those of the author(s) and do not necessarily reflect the views of the Department of the Treasury, the Mississippi Department of Environmental Quality, or the Mississippi Based RESTORE Act Center of Excellence.

